# A Lymphomimetic Synthetic Immune Niche, Consisting of CCL21 and ICAM1, Accelerates the Expansion of Potent CAR T-cells

**DOI:** 10.64898/2026.08.30.748076

**Authors:** Karin Brezinger-Dayan, Sofi Yado, Rawan Zoabi, Benjamin Geiger, Michal J Besser

## Abstract

**Background:** Chimeric Antigen Receptor (CAR) T-cell therapy has transformed the treatment of hematologic malignancies, yet, its broader clinical application often faces major challenges, including slow expansion rates, variable transduction efficiency, exhaustion, and functional heterogeneity. Recent studies have demonstrated that a Synthetic Immune Niche (SIN) composed of immobilized CCL21 and ICAM1 promotes the proliferation of murine and human T-cells while preserving their cytotoxic potency. In this study, we explored the capacity of immobilized CCL21 and ICAM1 to enhance the production of highly potent CAR T-cells by facilitating both their expansion and cytotoxic capacity.

**Methods:** CD19-directed CAR T-cells were generated from PBMCs of healthy donors using a clinical-grade protocol. Following retroviral transduction, the cells were expanded on CCL21 and ICAM1 coated plates, or on uncoated control plates. The effects of the SIN treatment on CAR T-cell expansion, morphology, physical properties, phenotypic markers, and potency were systematically evaluated.

**Results:** SIN exposure significantly enhanced CAR T-cell expansion, achieving a 9.3-fold higher total cell yield by day 13, compared with control cultures. This proliferative advantage persisted even after withdrawal of the cells from the synthetic niche on day 10. SIN stimulation preferentially expanded the CAR-transduced population, increasing CAR T-cell frequencies from 45.4% to 77.6%, resulting in a 14.6-fold increase in the absolute number of CAR T-cells compared with untreated cultures. Morphological and phenotypic analyses revealed distinct activated cell morphology, manifested by increased cell size, polarity and granularity, and elevated expression of the activation markers CD137 and CD69. Importantly, CAR T-cells transiently exposed to the SIN retained cytokine secretion and cytotoxic activity comparable to that of continuously SIN-treated cells, indicating that the SIN effect is persistent. Overall, SIN-conditioned CAR T-cells displayed robust antigen-dependent IFN-γ secretion and cytotoxicity against CD19-expressing target cells.

**Conclusion:** Stimulation of CAR T-cells with a synthetic immune niche consisting of immobilized CCL21 and ICAM1 enhances overall expansion while selectively enriching the CAR-transduced population, thereby substantially increasing both the number and prominence of therapeutically relevant CAR T-cells. This effect is accompanied by a persistent activation phenotype and high functional potency. We propose that incorporating SIN stimulation into CAR T-cell manufacturing represents a simple and scalable strategy for improving CAR T-cell yield while maintaining high cytotoxic efficacy.

## Introduction

Adoptive T-cell therapy (ACT) has become a transformative approach to cancer treatment, offering a potential for highly personalized and targeted therapies ^1–4^. Among its various strategies, Chimeric Antigen Receptor (CAR) T-cell therapy has achieved remarkable clinical success in hematologic malignancies ^5^. CAR T-cells are polyclonal T lymphocytes isolated from the patient’s peripheral blood and genetically engineered to express a CAR construct that confers specificity toward a defined tumor-associated antigen and incorporates intracellular co-stimulatory signaling domains, thereby enabling antigen recognition, independently of major histocompatibility complex (MHC) presentation ^5^. Clinically, CAR T-cell therapies are currently approved by the Food and Drug Administration for multiple B-cell malignancies, including acute lymphoblastic leukemia (ALL), diffuse large B-cell lymphoma (DLBCL), and multiple myeloma ^6^.

While CAR T-cell therapy offers unique advantages, its broader clinical application remains limited by several major challenges. These include variable transduction efficiency, limited persistence, and functional exhaustion, which collectively continue to constrain the efficacy and reproducibility across patients ^7^. Beyond CAR construct design, accumulating evidence indicates that *ex vivo* manufacturing conditions critically shape CAR T-cell fitness, influencing the expansion capacity, differentiation state, and cytotoxic potency ^5–7^. From a quantitative perspective, effective CAR T-cell-based therapies typically require cell numbers in the order of 10^8^–10^9^. However, at this expansion scale, T-cells frequently exhibit reduced cytotoxic potency, a phenomenon commonly associated with exhaustion, anergy, and increased susceptibility to inhibitory signals ^8,9^. These challenges are closely linked to the nature of current *ex vivo* activation and expansion protocols. In this context, conventional CAR T-cell manufacturing typically relies on strong stimulation in suspension culture, commonly using anti-CD3 or anti-CD3/CD28 activation platforms together with soluble cytokines, which can promote differentiation, functional heterogeneity, and progressive dysfunction during prolonged culturing protocols ^8,9^.

To address these broader challenges in CAR T-cell manufacturing, we recently developed 2D functionalized surfaces that act as a “Synthetic Immune Niche” (SIN) that supports the *ex vivo* expansion of T-cells while maintaining their intrinsic cytotoxic activity. This SIN consists of a tissue culture surface coated with the chemokine C-C motif Ligand 21 (CCL21) and the Intercellular Adhesion Molecule 1 (ICAM1) ^10,11^. CCL21, produced by stromal and endothelial cells within the lymph node ^12^, serves as a key regulator of immune responses, including the recruitment of T-cells and dendritic cells ^13,14^ guidance of immune cell migration ^15^, priming of T-cells for immune synapse formation ^16^ and co-stimulation of naïve T-cell expansion and Th1 polarization ^17,18,19^. CCL21 exerts these effects through its receptor CCR7, which is highly expressed on naïve and central memory T cells and plays a key role in lymph node homing, migration, and T-cell priming ^20^. ICAM1 is a key adhesion molecule that promotes immune synapse formation and T-cell activation through its interaction with the integrin receptor LFA- 1 (lymphocyte function–associated antigen 1) ^21^. Importantly, these two factors act synergistically, both *in vivo* and *ex vivo*, as CCL21 signaling enhances LFA-1 responsiveness to ICAM1 and mediates an arrest of motile lymphocytes on ICAM1-expressing dendritic cells, endothelial cells, and neighboring T-cells. This coordinated interaction facilitates efficient T-cell priming and activation within the physiological immune microenvironment ^22,23^.

In our previous studies, we demonstrated that culturing activated murine CD4^+^ ^10^ or CD8^+11^ T-cells on the immobilized CCL21+ICAM-based SIN markedly enhanced the expansion of both T-cell subsets. Furthermore, treatment of ovalbumin (OVA)-specific CD8⁺ T-cells with this SIN increased their cytotoxic activity against OVA-expressing B16 melanoma target cells, as well as their tumor suppressive activity in vivo ^11^. In addition, a recent study demonstrated that SIN stimulation differentially modulates the balance between proliferation and cytotoxicity in an activation mode-specific manner ^24^. At the molecular level, this differential SIN-mediated regulation was associated with specific molecular programs linked to proliferation, cytotoxic effector function and exhaustion, with SIN-treated cells exhibiting reduced expression of exhaustion-associated genes ^24^. Collectively, these results highlight the capacity of the CCL21+ICAM1-based SIN to induce an “optimal interplay” between the proliferation and cytotoxicity of CD8^+^ T-cells, supporting its potential to reinforce adoptive cancer immunotherapy ^24^.

More recently, we extended these findings to a clinically relevant setting and showed that the CCL21+ICAM1 SIN affects patient-derived tumor-infiltrating lymphocytes (TILs) behavior, influencing activation-associated features and enhancing both expansion and cytotoxic readouts in a donor-specific manner, thereby supporting the potential of SIN to optimize the clinical utility of adoptive T-cell products ^25^. Notably, application of the CCL21+ICAM1-based SIN to patient-derived TILs revealed substantial inter-patient variability in expansion outcomes during the rapid expansion protocol (REP) ^25^. Subsequent morphological and molecular profiling of pre-REP TILs enabled prediction of intrinsic proliferative potential, enabling the development of tailored SIN-based strategies that differentially enhance the expansion and cytotoxic function across TIL subsets ^26^. Despite this growing body of work across diverse T-cell settings, including patient-derived TILs, the impact of SIN-mediated microenvironmental cues on CAR T-cells remains largely unexplored. Importantly, CAR T-cell immune synapses have been shown to be structurally and functionally distinct from classical TCR synapses, raising the question of whether lympho-mimetic cues such as those induced by the immobilized CCL21 and ICAM1, can beneficially reprogram CAR T-cell expansion and effector function during their production ^27^.

In the present study, we applied the CCL21+ICAM1-based SIN technology to human CD19- directed CAR T-cells to systematically evaluate its effects on CAR T-cell expansion, phenotype, functional potency, and antitumor activity. We demonstrate here, for the first time, that SIN stimulation markedly enhances CAR T-cell expansion, selectively enriches the CAR-transduced population, induces a sustained activation state characterized by distinct morphological and phenotypic changes, while preserving antigen-specific cytotoxic function. Moreover, transient exposure to the SIN was sufficient to induce a durable priming effect that persisted also after removal of the cells from the synthetic niche. These findings identify this lympho- mimetic microenvironmental conditioning as a simple and scalable strategy for improving both the yield and composition of CAR T-cell products without compromising functional potency.

## Materials and Methods

### Cell Lines and Culture Conditions

The CD19-expressing human B-cell acute lymphoblastic leukemia (B-ALL) cell line NALM-6 and the CD19-negative human multiple myeloma cell line U266 were obtained from ATCC and cultured in RPMI 1640 medium (Gibco, Thermo Fisher Scientific, Waltham, MA, USA), supplemented with 10% fetal bovine serum (FBS), 2mM L-glutamine, 1% penicillin-streptomycin (all from Gibco) and 1 mM sodium pyruvate (Sartorius, Göttingen, Germany), hereafter referred to as complete medium (CM). Cells were cultured at 37 °C in a humidified atmosphere containing 5% CO₂, passaged every 48-72 h. Cells in the logarithmic growth phase were used for all experiments. NALM-6 and U266 cells served as CD19-positive and CD19- negative target cells, respectively, in cytotoxicity assays, as described below.

### Primary Human Samples and Study Approval

Peripheral blood mononuclear cells (PBMCs) were isolated from three healthy, unrelated donors, using a density gradient centrifugation method with Ficoll-Hypaque (Lymphoprep, Serumwerk Bernburg AG, Germany). Written informed consent was obtained from all donors, and all procedures were approved by the Institutional Review Board (IRB) of the Rabin Medical Center (approval no. 0428-25-RMC).

### Substrate Functionalization

Substrate functionalization was performed by overnight incubation in phosphate-buffered saline (PBS, Sartorius) with 10 µg/mL hCCL21 and 100 µg/mL hICAM1 (produced by the Structural Proteomics Unit, Weizmann Institute of Science, Rehovot, Israel). Detailed information on protein production and purification can be found in the Supplementary Methods section.

### T-cell Activation and Transduction

On the day of culture initiation (Day 0), PBMCs were thawed and incubated in CM supplemented with 300 IU/mL recombinant human interleukin-2 (IL-2; Aldesleukin; Iovance, Basel, Switzerland). T-cell activation was initiated by the addition of anti-CD3 monoclonal antibody (OKT-3; 50 ng/mL; Miltenyi Biotec, Bergisch Gladbach, Germany). Following 48 h of stimulation (Day 2), the activated cells were transduced with a third-generation anti-CD19 retroviral vector (RVV), kindly provided by Brenner MK, Baylor College of Medicine. For transduction, non–tissue culture treated 6-well plates were coated overnight at 4 °C with RetroNectin (10 µg/mL; Takara Bio Inc., Otsu, Japan) in PBS (Sartorius). Retroviral supernatant was rapidly thawed and diluted 1:4 in RPMI 1640 medium supplemented with 10% FBS. Five milliliters of diluted vector were added per well, followed by centrifugation at 2,000 × g for 1 h at 32°C. The supernatant was then aspirated, and cells were resuspended in CM supplemented with 300 IU/mL IL-2. A total of 2.5 × 10⁶ cells were seeded per well, centrifuged at 1,000 × g for 10 min, and incubated overnight at 37 °C with 5% CO₂.

### Stimulation of CAR T-cells with Surface-Immobilized CCL21 and ICAM1 SIN

On Day 3, transduced cells were washed and transferred to either CCL21+ICAM1-coated 24- well plates or standard non-coated control plates at a density of 1 × 10⁵ cells per well in 1 mL of CM supplemented with 300 IU/mL IL-2. Freshly CCL21+ICAM1-coated plates were prepared one day prior to each cell seeding or cell splitting. Cells were cultured for a total of 13 days and split on days 7 and 10 to maintain a cell concentration of 0.1–2.5×10^5^ cells/mL. At each splitting point, cells were transferred to freshly coated plates to ensure continuous exposure to SIN stimulation. To determine whether the SIN-induced effects were maintained even in the absence of CCL21 and ICAM1, SIN-stimulated cells were transferred on day 10 to non-coated plates and cultured for an additional period of 72 h prior to analysis, while cells under continuous SIN exposure remained on coated plates until day 13.

The number of viable cells was determined on days 6, 7, 10, and 13 using an automated cell counter device (Countess II; Thermo Fisher Scientific) and trypan blue staining solution (Gibco). Fold expansion at each time point was calculated relative to the initial number of cells seeded on day 3.

### Spectral Flow Cytometry

CAR transduction efficiency was evaluated on days 6, 10 and 13 by spectral flow cytometry. Cells were stained with a biotin-conjugated Protein L antibody (GenScript, Piscataway, NJ, USA), followed by streptavidin-PE (Biolegend, San Diego, CA, USA), and co-stained with anti-human CD3 (APC; Miltenyi Biotec). CD3⁺Protein L⁺ cells were defined as transduced CAR T-cells. Untransduced cells served as negative controls.

To define molecular phenotypic profiles, surface marker expression was analyzed on days 6, 10, and 13 by spectral flow cytometry. At the indicated time points, the cells were detached from the substrate and washed with 5% FBS / 0.5 mM EDTA in PBS. The supernatant was aspirated, and the cells were surface stained at 4°C for 10 min with Zombie viability dye (BioLegend) to exclude dead cells and with the following fluorescent monoclonal antibodies: CD4-PE Vio760, CD8-APC Vio770, CCR7-Vio Bright V600, CD45RA-VioBlue, CD69-PE Vio770, CD137/4-1BB-Vio Bright R720, PD-1-Vio Bright V423 and TIM-3-Vio Bright FITC (all from Miltenyi Biotec). After washing, data were acquired using a Northen Lights (Cytek, CA, USA) spectral flow cytometer and analyzed using SpectroFlo software and FlowJo software (Ashland, OR, USA).

### Microscopy and Image Analysis

The morphology and growth patterns of the CAR T-cells were examined using phase-contrast microscopy (Olympus CKX53) with a 20x objective lens. Images were taken on days 6, 7, 10 and 13 and analyzed using ImageJ software (https://imagej.nih.gov/ij/). To quantify the projected area and circularity of CAR T-cells on day 9, we developed a model in Cellpose (v 3.1.0) ^28^, using the built-in “cyto2-cp3” model as the starting point followed by further training. The resulting masks were post-processed using the MorphoLibJ plugin ^29^,where boundary erosion was applied to remove halo and shadow artifacts and improve contour accuracy. Cells located at image borders were excluded from analysis. In addition, objects with an area <30 µm² were filtered out as likely debris.

### IFN-γ Secretion and Cell-Mediated Cytotoxicity

To evaluate antigen-specific functionality, IFN-γ secretion was measured following co-incubation of CAR T-cells, pre-cultured on non-coated substrates or CCL21+ICAM1-coated surfaces, with the CD19-positive NALM-6 cell line. The CD19-negative cell line U266 served as negative control. Co-cultures were performed in 96-well plates on days 6 and 13 at an effector-to-target (E:T) ratio of 1:1 (1 × 10⁵ cells each) in a total volume of 200 µL CM and incubated overnight at 37°C with 5% CO₂. Supernatants were collected, diluted as required, and IFN-γ levels were quantified by ELISA using the Human IFN-γ ELISA MAX Deluxe Set (BioLegend, San Diego, CA, USA), according to the manufacturer’s instructions.

Cytotoxic activity was assessed in parallel by measuring lactate dehydrogenase (LDH) release into the culture supernatant using a colorimetric assay (CyQUANT LDH Cytotoxicity Assay Kit, Thermo Fisher Scientific), according to the manufacturer’s instructions. Target cell viability was calculated from LDH release measurements and expressed as the percentage of viable cells relative to the maximum viability control. Viability (%) was determined as 100 minus the percentage of specific lysis, where specific lysis was calculated as [(experimental LDH release − spontaneous LDH release) / (maximum LDH release − spontaneous LDH release)] × 100. Spontaneous LDH release was defined as LDH released from target cells cultured alone, and maximum LDH release was determined by complete lysis of target cells using the kit-provided lysis buffer. Absorbance was measured at 490 nm using a microplate reader.

### Statistical Analysis

Statistical analyses were performed using two-tailed paired Student’s *t*-tests for pairwise comparisons between experimental groups. A *p*-value ≤ 0.05 was considered statistically significant (*p < 0.05, **p < 0.01, ***p < 0.001).

## Results

### CCL21+ICAM1-Based SIN Enhances CAR T-cell Proliferation

To evaluate the impact of a CCL21+ICAM1-based Synthetic Immune Niche (SIN) on CAR T-cell expansion, we utilized a clinical-grade manufacturing protocol. PBMCs from three healthy donors were stimulated with anti-CD3 and IL-2 (day 0) and subsequently transduced on day 2 with anti-CD19 retroviral vector (RVV). This RVV, currently used in our clinical trial, was selected for its moderate multiplicity of infection (MOI = 1.5), which typically yields a 50% to 60% transduction efficiency, a range optimal for observing the differential modulatory effects of the SIN on transduced versus untransduced cells. On day 3, cells were transferred to either standard tissue culture 24-well plates (“non-coated”) or plates coated with CCL21 and ICAM1 (“SIN-coated”). Cultures were maintained through day 13, with periodic splitting and re-exposure to freshly coated SIN substrates on days 7 and 10. To assess whether the proliferative advantage induced by the SIN required continuous niche exposure, a subset of CCL21+ICAM1-exposed cells was transferred to non-coated plates on day 10 for the final 72 hours of culture (“SIN to non-coated”). Fold expansion at each time point was calculated relative to the initial number of cells seeded on day 3.

Throughout the expansion period, cell viability remained consistently high (≥ 90%) across all donors and culture conditions, indicating that the CCL21+ICAM1-based SIN does not induce acute toxicity. As shown in **Figure 1A**, initial assessments at early time points (days 6 and 7) revealed a trend toward increased expansion in SIN-exposed cultures, which did not reach statistical significance (*p - values ≥* 0.479). However, a pronounced proliferative benefit emerged at later time points. By day 10, SIN-exposed cells showed significantly higher average fold expansion values (mean ± SD, 154.3 ± 5.9) compared to untreated cells (73.0 ± 41.0, *p* = 0.0499) **(Figure 1B)**. This effect was further amplified by day 13, with SIN-exposed cells exhibiting significantly greater expansion (910.0 ± 53.5 vs. 209.5 ±170.7 in untreated cells, *p* = 0.0052) **(Figure 1B)**.

**Figure 1.**
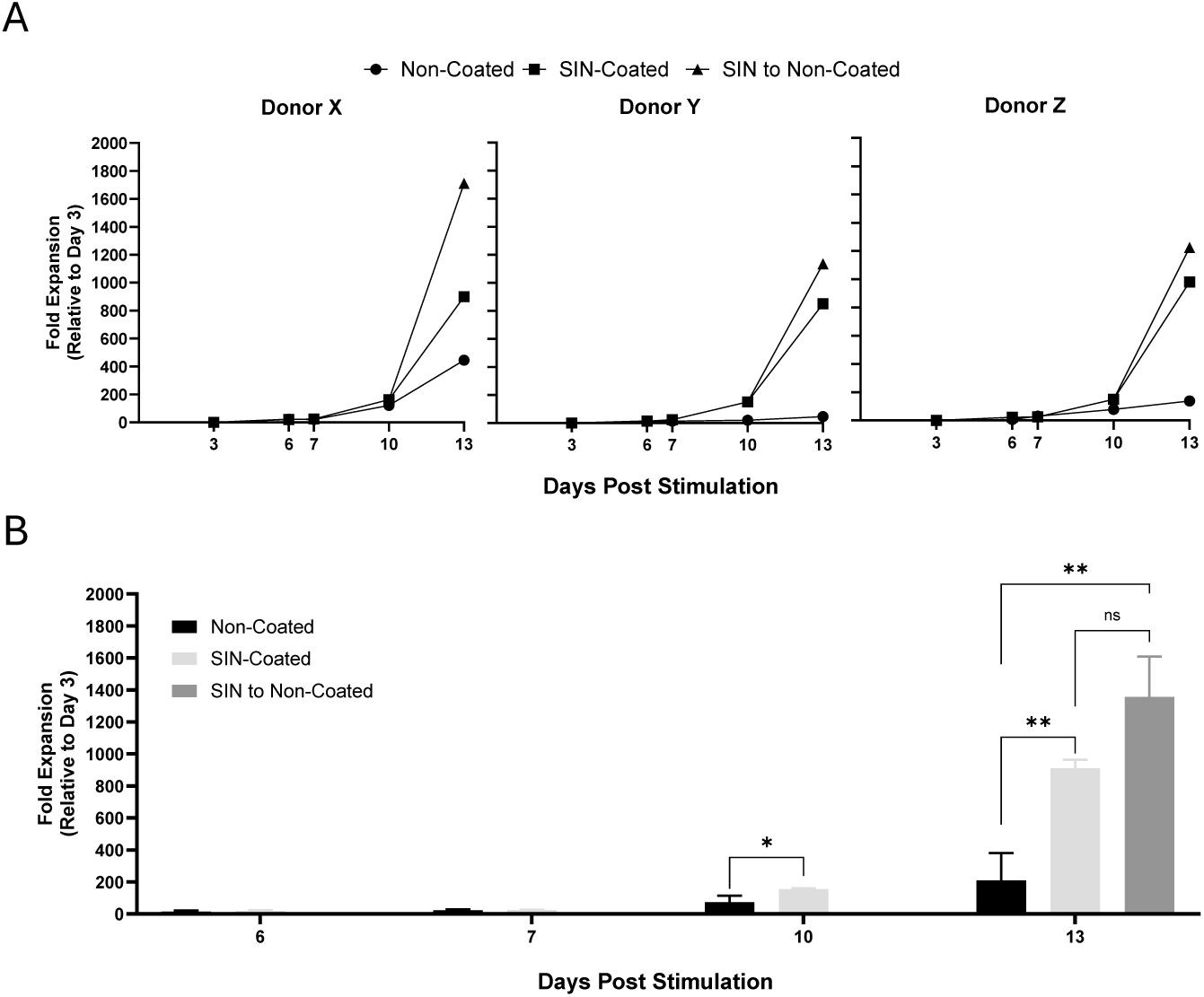
CCL21+ICAM1 exposure enhances the CAR T-cell proliferation over time. Results are based on CAR T-cells prepared from 3 independent donors (X, Y, Z) and examined for 13 days. One sample of each donor (“SIN to non-coated”) was SIN-stimulated for 10 days and further cultured in non-coated plates. **(A)** Fold expansion of individual CAR-T cultures. **(B)** Average fold expansion of all three donors. *p <.05, **p <.01.

Analysis of donor-specific responses revealed a mean 9.3-fold increase in expansion relative to untreated cells (range, 2.0-18.6), indicating that although the magnitude of the response varied among donors, all donors exhibited enhanced expansion following SIN stimulation. Importantly, this proliferative effect was sustained even after removing the cells from the synthetic niche. Specifically, CAR T-cells that expanded on the SIN-coated surfaces until day 10 and subsequently transferred to non-coated plates retained significantly greater expansion by day 13 than untreated cultures (1357.0 ± 252.5 vs. 73.0 ± 41.0, *p* = 0.006) **(Figure 1B)**.

Collectively, these data demonstrate that exposure to a CCL21+ICAM1-functionalized surface significantly enhances the proliferative capacity of CAR T-cells without compromising viability and further suggest that this pro-proliferative "priming" effect persists for at least 3 days, even after the cells are removed from the synthetic niche.

### SIN exposure induces morphological transformation in CAR T-cells, manifested by large size and polarized morphology

The enhanced proliferative capacity of SIN-treated CAR T-cells was accompanied by distinct morphological changes. Microscopy-based imaging revealed that CAR T-cells, expanded under standard, “non-coated” conditions, remained mostly spherical (median circularity, IQR: 0.926, [0.909-0.933]) and predominantly small (median projected area:, IQR: 61.47, [51.83- 75.12] µm²), whereas SIN-exposed cultures displayed a high frequency of polarized (median circularity, IQR: 0.772, [0.637-0.882]), elongated, or "crescent-shaped" larger cells (median projected area:,IQR: 112.12, [80.78-144.69] µm², **Supplementary Figure 1a-c**). These morphological differences were consistently observed across all donors on day 9 **(Figure 2A)**, confirming the reproducibility of the observed phenotype. The same pattern was also observed on day 13, as shown for donor x in **Figure 2B**. Such morphological asymmetry is a hallmark of T-cell activation and cytoskeletal remodeling processes ^30–33^, suggesting a more dynamic and functionally activated cellular phenotype.

**Figure 2.**
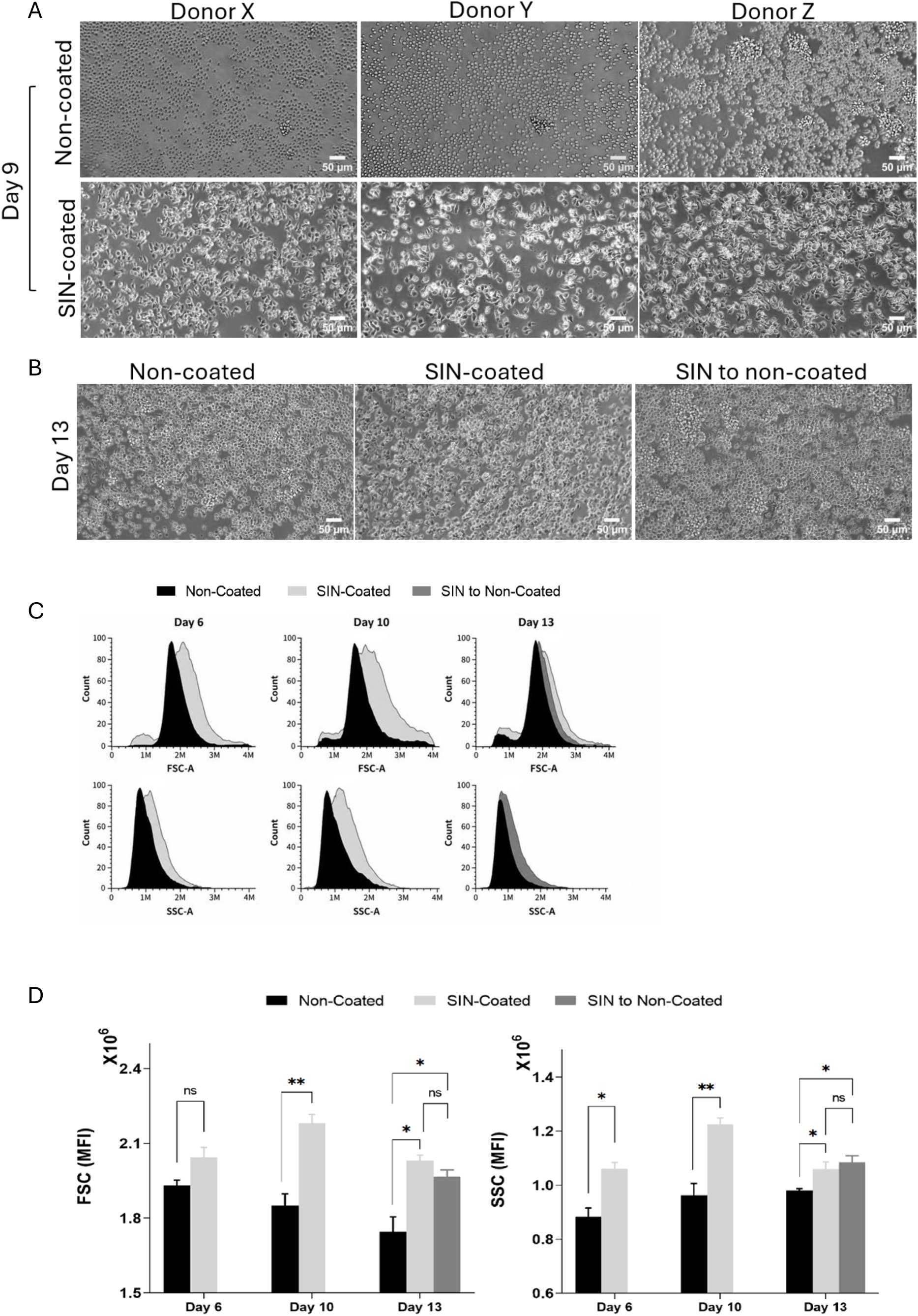
SIN exposure induces morphological transformation in CAR T-cells, manifested by large size and polarized morphology. **(A)** Phase contest microscopy images of CAR T-cell cultures from all donors on day 9. Scale bar, 50µm. **(B)** Representative microscopy images from donor x on day 13 of SIN-treated cultures, untreated cultures, and cells initially exposed to SIN and transferred to non-coated plates on day 10. Scale bar, 50µm. **(C)** Representative flow cytometry histograms of FSC and SSC from donor x on days 6, 10 and 13. The SSC histograms of day 13 from SIN-coated and SIN to non-coated completely overlap. **(D)** Quantitative analysis of FSC (left) and SSC (right) across all three donors on days 6, 10, and 13. *p <.05, **p <.01.

Quantitative flow cytometry analysis further corroborated these findings, showing that SIN-stimulated CAR T-cells displayed significantly higher Forward Scatter (FSC) values on days 10 (*p* = 0.0048) and 13 (*p* = 0.0107), as well as increased Side Scatter (SSC) intensities across all analyzed time points (day 6, *p* = 0.0105; day 10, *p* = 0.006; day 13, *p* = 0.0473), compared to untreated cells. Representative flow cytometry histograms from donor X are shown in **Figure 2C**, and the quantitative analysis of all three donors is presented in **Figure 2D**. The increased FSC and SSC are consistent with increased cell size and greater cellular granularity or structural complexity, respectively, features associated with T-cell activation and blast transformation ^34^. Importantly, following SIN removal on day 10, CAR T-cells retained the enlarged and polarized morphology comparable to that of cells that were continuously SIN-treated **(Figure 2D)**. Taken together, these findings indicate that SIN induces distinct morphological and structural changes in CAR T-cells that are persistent for at least three days beyond direct SIN stimulation.

### SIN Treatment Preferentially Enhances the Proliferation of Transduced CAR-T Cells, thereby Increasing Their Prominence

Flow cytometry analysis conducted on days 6, 10, and 13 indicated that SIN treatment significantly enriched the frequency of CAR expressing cells within the culture. As early as day 6, CCL21+ICAM1-coated plates yielded a significantly higher percentage of CAR T-cells (46.8 ± 3.2%) compared to non-coated controls (39.5 ± 0.5%, *p* = 0.032). This difference increased substantially as the culture progressed; by day 10, SIN-exposed cultures reached 59.3 ± 4.4% CAR T-cells (vs. 34.7 ± 0.7%, *p* ≤ 0.001), and by day 13, this enrichment further increased to 77.6 ± 5.0% (vs. 45.4 ± 6.1%, *p* = 0.004) **(Figure 3A** **and** **Table 1)**. In contrast, a stable frequency of CAR T-cells throughout the expansion phase was observed during GMP manufacturing of clinical CD19 CAR T-cell products **(Supplementary Figure S2)**. Because the total cell numbers, overall fold expansion, and the frequency of transduced CAR T-cells were known for each culture condition, we next determined the absolute number of CAR positive T-cells at each time point by multiplying the total cell count by the corresponding frequency of CAR T-cells. As shown in **Figure 3C**, the preferential enrichment of CAR T-cells observed in SIN-treated cultures translated into a striking increase in the absolute number of CAR T-cells. By day 13, CAR T-cells cultured continuously on SIN-coated plates exhibited a 14.6-fold greater expansion (range 4.3-28.5, p = 0.0002) than CAR T-cells cultured on non-coated plates. CAR T-cells exposed to SIN until day 10 and subsequently transferred to uncoated plates demonstrated an even greater 19.4-fold expansion advantage over untreated controls (range 7.9-37.3, p = 0.0046). In contrast, neither the absolute number nor the fold expansion of the non-transduced CAR negative cell population was significantly affected by exposure to the CCL21+ICAM1-coated surfaces at any time point (all p ≥ 0.118) **(Figure 3D)**.

**Figure 3.**
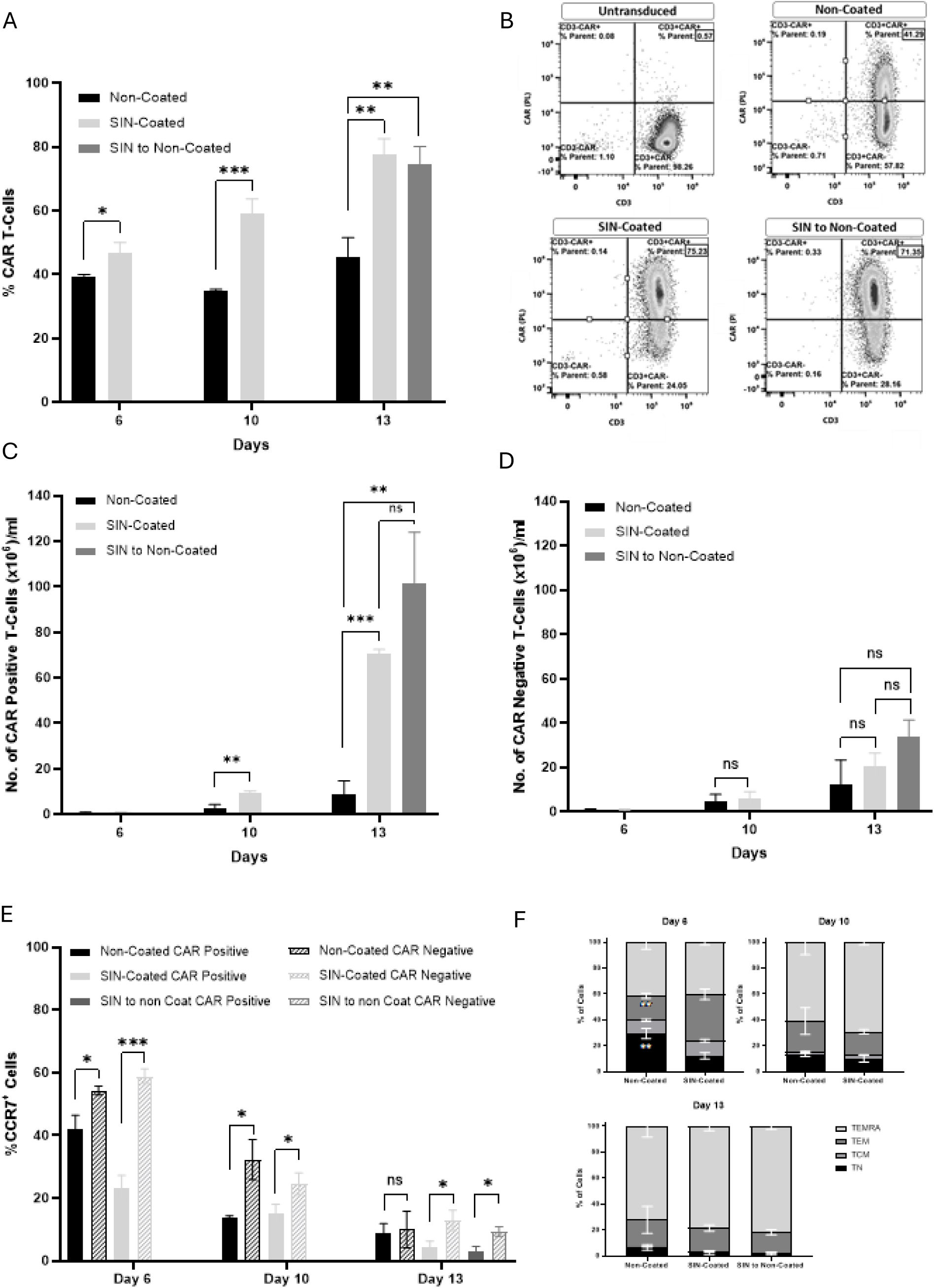
SIN-coated plates enhance CAR-T-cell expansion and promote early differentiation. **(A)** Frequency of transduced CAR T-cells determined by labeling for Protein L⁺/CD3⁺ during culture; **(B)** Representative flow cytometry dot plots showing Protein L⁺/CD3⁺ CAR T-cells on day 13; **(C)** Absolute number of CAR-positive T-cells; **(D)** absolute number of CAR-negative T-cells; **(E)** Percentage of CCR7⁺ cells among CAR-positive and CAR-negative T-cell populations during culture; **(F)** Distribution of CAR T-cell differentiation subsets, including naive T cells (TN; CD45RA⁺CCR7⁺), central memory T cells (TCM; CD45RA⁻CCR7⁺), effector memory T cells (TEM; CD45RA⁻CCR7⁻), and terminally differentiated effector memory T cells (TEMRA; CD45RA⁺CCR7⁻). Data represent mean ± SD (n = 3); *p < .05, **p < .01, ***p < .001; ns, not significant.

**Table 1.** Phenotypic comparison of non-coated and SIN-coated CAR T-cells from three donors shown as average ± SD.

| Category | Parameter | Condition | Non-coated vs. SIN-coated |  |  | SIN vs. SIN to non-coat |
| --- | --- | --- | --- | --- | --- | --- |
|  |  |  | Day 6 | Day 10 | Day 13 | Day 13 |
| CAR T-cell | % CD3<sup>+</sup> ProteinL<sup>+</sup> | Non-coated | 39.5 ± 0.5 | 34.7 ± 0.7 | 45.4 ± 6.1 | |
| | | SIN-coated | 46.8 ± 3.2 | 59.3 ± 4.4 | 77.6 ± 5.0 | 77.6 ± 5.0 |
| | | SIN to non-coat | | | | 74.7 ± 5.4 |
|  |  | p-value | **0.032** | **≤ 0.001** | **0.004** | 0.609 |
| T-cell subpop. | % CD8<sup>+</sup> | Non-coated | 63.2 ± 4.6 | 74.02 ± 3.9 | 80.0 ± 3.8 | |
| | | SIN-coated | 59.0 ± 6.6 | 74.1 ± 5.7 | 86.5 ± 3.1 | 86.5 ± 3.1 |
| | | SIN to non-coat | | | | 88.3 ± 3.1 |
|  |  | p-value | 0.497 | 0.981 | 0.135 | 0.585 |
| | % CD4<sup>+</sup> | Non-coated | 35.8 ± 4.6 | 25.3 ± 3.9 | 18.5 ± 4.6 | |
| | | SIN-coated | 39.5 ± 6.5 | 25.2 ± 5.7 | 12.8 ± 3.1 | 12.8 ± 3.1 |
| | | SIN to non-coat | | | | 10 ± 3.1 |
|  |  | p-value | 0.543 | 0.986 | 0.218 | 0.536 |
| Activation Markers | % CD137<sup>+</sup> | Non-coated | 2.1 ± 0.6 | 0.9 ± 0.1 | 1.3 ± 0.5 | |
| | | SIN-coated | 45.1 ± 7.4 | 23.3 ± 5.4 | 10.8 ± 3.4 | 10.8 ± 3.4 |
| | | SIN to non-coat | | | | 10.6 ± 5.1 |
|  |  | p-value | **0.001** | **0.004** | **0.018** | 0.975 |
| | % CD69<sup>+</sup> | Non-coated | 9.7 ± 2.5 | 12.0 ± 2.8 | 24.2 ± 6.0 | |
| | | SIN-coated | 18.0 ± 3.5 | 31.6 ± 7.2 | 31.4 ± 7.5 | 31.4 ± 7.5 |
| | | SIN to non-coat | | | | 31.5 ± 9.8 |
|  |  | p-value | 0.051 | **0.023** | 0.352 | 0.991 |
| | % CD137<sup>+</sup>CD69<sup>+</sup> | Non-coated | 0.6 ± 0.2 | 0.6 ± 0.02 | 1.8 ± 0.7 | |
| | | SIN-coated | 12.1 ± 3.4 | 12.7 ± 3.4 | 6.3 ± 1.8 | 6.3 ± 1.8 |
| | | SIN to non-coat | | | | 7.3 ± 3.3 |
|  |  | p-value | **0.01** | **0.008** | **0.028** | 0.736 |
| Activation associated checkpoints | % PD-1<sup>+</sup> | Non-coated | 0.2 ± 0.1 | 0.3 ± 0.1 | 0.6 ± 0.2 | |
| | | SIN-coated | 8.9 ± 2.5 | 7.7 ± 1.00 | 1.8 ± 0.5 | 1.8 ± 0.5 |
| | | | | | | 1.2 ± 0.2 |
|  |  | p-value | **0.01** | **0.001** | **0.024** | 0.063 |
| | % TIM-3<sup>+</sup> | Non-coated | 89.9 ± 3.6 | 59.9 ± 10.2 | 68.4 ± 9.5 | |
| | | SIN-coated | 91.8 ± 3.2 | 77.8 ± 4.8 | 84.8 ± 7.3 | 84.8 ± 7.3 |
| | | | | | | 79.8 ± 10.3 |
|  |  | p-value | 0.61 | **0.087** | 0.12 | 0.308 |
| | % PD-1<sup>+</sup>TIM-3<sup>+</sup> | Non-coated | 0.2 ± 0.1 | 0.3 ± 0.2 | 0.8 ± 0.2 | |
| | | SIN-coated | 16.1 ± 4.9 | 5.0 ± 0.8 | 2.9 ± 0.8 | 2.9 ± 0.8 |
| | | | | | | 0.6 ± 0.2 |
|  |  | p-value | **0.01** | **0.001** | **0.03** | **0.019** |
| Differentiation status | Naive | Non-coated | 29.5 ± 3.9 | 12.9 ± 1.2 | 6.6 ± 2.2 | |
| | | SIN-coated | 12.3 ± 2.3 | 10.2 ± 2.7 | 2.7 ± 1.5 | 2.7 ± 1.5 |
| | | | | | | 1.7 ± 1.1 |
|  |  | p-value | **0.006** | 0.255 | 0.105 | 0.491 |
| | TCM | Non-coated | 10.3 ± 0.9 | 2.5 ± 0.4 | 0.5 ± 0.1 | |
| | | SIN-coated | 11.7 ± 1.0 | 2.7 ± 0.1 | 0.9 ± 0.4 | 0.9 ± 0.4 |
| | | | | | | 0.8 ± 0.3 |
|  |  | p-value | 0.207 | **0.381** | 0.272 | 0.793 |
| | TEM | Non-coated | 19.0 ± 1.9 | 23.7 ± 10.4 | 20.6 ± 10.4 | |
| | | SIN-coated | 35.8 ± 4.1 | 17.9 ± 1.8 | 17.9 ± 2.4 | 17.9 ± 2.4 |
| | | | | | | 15.8 ± 2.1 |
|  |  | p-value | **0.007** | 0.479 | 0.734 | 0.401 |
| | TEMRA | Non-coated | 41.2 ± 5.3 | 60.9 ± 9.6 | 72.2 ± 8.2 | |
| | | SIN-coated | 40.2 ± 2.0 | 69.2 ± 1.9 | 78.5 ± 3.4 | 78.5 ± 3.4 |
| | | | | | | 81.7 ± 2.4 |
|  |  | p-value | 0.819 | **0.296** | 0.375 | 0.338 |

These findings indicate that the proliferative effect of the SIN is not uniformly distributed across the culture but is preferentially directed toward the CAR-transduced population, thereby substantially increasing both the yield, and thereby the relative prominence of the therapeutically relevant CAR T-cells.

To explore potential mechanisms underlying the preferential expansion of the CAR-transduced population, we compared the expression level of CCR7, the receptor for CCL21, on CAR-positive and CAR-negative cells throughout the culture period. As shown in **Figure 3E**, CCR7 expression progressively declined over time in both populations. However, at each time point, CAR-negative cells consistently expressed significantly higher levels of CCR7 than CAR-positive cells within the same culture (all p ≤ 0.0428). These findings indicate that preferential expansion of the CAR-transduced population is not associated with an increased CCR7 expression.

We next characterized the differentiation and activation profile of the CAR T-cell cultures. SIN coating did not significantly alter the overall CD4/CD8 ratio, either within the total culture or within the CAR T-cell subpopulation (p ≥ 0.218) **(Table 1)**. On day 6, SIN-expanded CAR T-cells showed an accelerated shift from a naive phenotype (TN; CCR7^+^CD45RA^+^) toward an effector memory phenotype (TEM; CCR7^-^CD45RA^-^), with TEM frequencies significantly higher in SIN-stimulated cultures (35.8 ± 4.1%) than in controls (19.0 ± 1.9%, *p* = 0.007; **Figure 3F**). These early phenotypic differences were no longer evident on days 10 and 13, at which point the majority of CAR T-cells in all conditions had matured into terminally differentiated effector memory (TEMRA; CCR7^-^CD45RA^+^) cells **(Figure 3F** **and** **Table 1)**, suggesting that SIN exposure accelerates early differentiation dynamics without altering the final differentiation endpoint.

Analysis of activation markers revealed a marked increase in CD137 (4-1BB) and CD69 expression in SIN-exposed cells **(Table1)**. While CD137 expression remained minimal in cells growing in non-coated plates (0.9-2.1%), SIN-stimulated cells exhibited substantially higher levels, which gradually declined over time (day 6: 45.1 ± 7.4%; day 10: 23.3 ± 5.4%; day 13: 10.7 ± 3.4%; *p*-values < 0.02; **Table 1**). Similarly, CD69 expression was consistently higher in SIN-stimulated cells, reaching statistical significance on day 10 (31.6 ± 7.2% vs. 12.0 ± 2.8%, *p* = 0.023; **Table 1**). Accordingly, SIN-treated cultures contained significantly higher proportions -of CD137^+^CD69^+^ "strongly activated" CAR T-cells throughout the expansion period (p ≤ 0.028; **Table 1**).

In contrast, expression of the exhaustion-associated marker TIM-3 was high (60-91%) across all conditions and time points, independent of SIN exposure **(Table 1)**. The activation-associated checkpoint molecule PD-1 remained generally low (≤ 8.9%), although modest but significant increases were observed in SIN-treated CAR T-cells compared to the near-zero levels in untreated cells (e.g., day 10: 7.7 ± 1.0% vs. 0.3 ± 0.1%, *p* < 0.001; **Table 1**). Notably, similar phenotypic profiles were observed in CAR T-cells maintained on SIN-coated surfaces through day 13 and in those transferred to non-coated conditions on day 10, indicating that SIN-induced activation states are maintained following niche withdrawal **(Table1)**. Collectively, these findings indicate that the CCL21+ICAM1-based SIN promotes enrichment of CAR T-cells and induces a robust and sustained activation state without evidence of excessive exhaustion.

### The Synthetic Immune Niche Preserves Antigen-Specific Cytotoxicity and Polyfunctional Potency

Finally, we directly evaluated the anti-tumor activity of the SIN-treated and non-treated CD19 CAR T-cells by co-culturing them with CD19⁺ NALM-6 leukemia cells, or with CD19⁻ U266 cells that served as negative controls. The CAR T-cells used for the cytotoxic assay were cultured for 6 or 13 days with SIN or non-coated plates, as well as with SIN for 10 days, followed by 3 days in non-coated plate (“SIN-to-non-coated cultures”).

We assessed polyfunctional potency of the CAR T-cells, and their ability to simultaneously mediate direct cytolysis and coordinate an immune response through cytokine secretion. As shown in **Figure 4A**, CAR T-cells from all three culture conditions exhibited strong antigen-specific cytokine responses, with markedly elevated IFN-γ secretion upon co-culture with CD19⁺ NALM-6 cells (108,020 ± 49,192 pg/ml) compared to CD19⁻ U266 cells (2,442 ± 1,943 pg/ml; p = 0.002). Importantly, upon exposure to NALM-6 targets, all CAR-T cells secreted high levels of IFN-γ, with no significant differences observed between non-coated, SIN-coated, and SIN-to-non-coated cultures (all p ≥ 0.280) **(Figure 4A)**.

**Figure 4.**
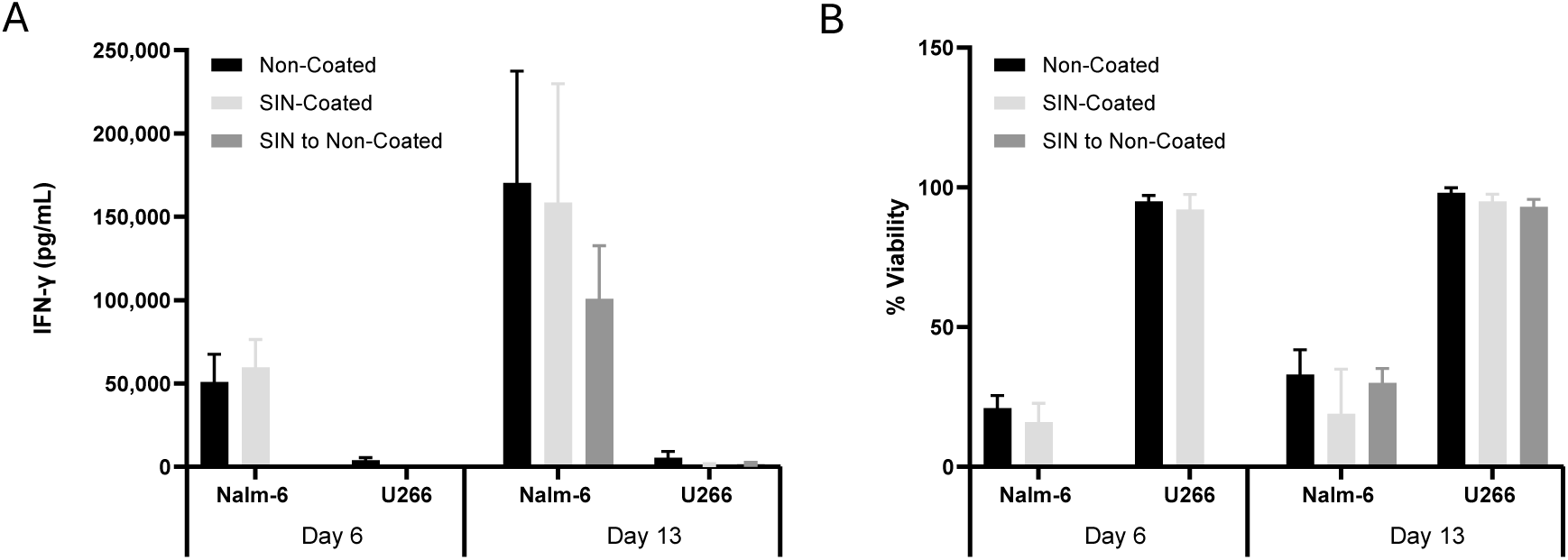
Functional assessment of SIN-expanded CAR T-cells. **(A)** IFNγ secretion **(B)** viability of CD19⁺ NALM-6 and CD19⁻ U266 target cells following overnight co-culture with CAR T-cells, showing persistent cytokine secretion and tumor-killing capacity across all conditions. Data represents mean ± SD (n = 3).

Antigen-specific cytotoxicity was further assessed by measuring target cell viability using an LDH-based assay. Co-culture with CAR T-cells resulted in a substantial reduction in NALM-6 cell viability (18.5 ± 4.5% on day 6 and 27.3 ± 8.9% on day 13), whereas U266 cells maintained high viability (>92%) across all conditions. Consistent with the cytokine data, no significant differences in cytotoxic activity were observed among non-coated, SIN-coated, and SIN-to- non-coated CAR T-cells (all p ≥ 0.357; **Figure 4B**).

Together, these results demonstrate that SIN stimulation enhances CAR T-cell expansion and activation while preserving key effector functions. Despite their markedly enhanced expansion and activated phenotype, CAR T-cells expanded under SIN conditions retain antigen specificity, robust cytokine secretion, and potent cytolytic activity against CD19⁺ tumor targets, indicating that SIN-induced phenotypic and proliferative changes do not compromise functional potency.

## Discussion

Adoptive cellular immunotherapies, including CAR T cells and tumor-infiltrating lymphocytes (TIL), have transformed the treatment landscape of hematologic and solid malignancies, yet their broader clinical implementation remains constrained by different manufacturing challenges ^1–4^. These include limited expansion capacity, prolonged culture duration, phenotypic heterogeneity, and declining cytotoxic potency during the ex *vivo* expansion. In addition, CAR T-cells often display considerable cellular heterogeneity due to variable transduction efficiency. In the present study, we demonstrate that the lympho-mimetic synthetic immune niche (SIN) composed of immobilized CCL21 and ICAM1 favorably addresses several of these limitations. Exposure of CD19-directed CAR T-cells to this SIN resulted in markedly enhanced expansion of the cells, a major increase in the prominence of the CAR-transduced cell population, sustained activation, and preservation of antigen-specific cytotoxic functionality.

One of the most striking findings reported here is the pronounced proliferative advantage conferred by SIN stimulation. By day 13, cells cultured on SIN-coated surfaces achieved a 9.3- fold greater expansion than cells cultured under standard conditions. Importantly, this effect persisted after removal of the cells from the niche. CAR T-cells transferred from SIN-coated to non-coated plates on day 10 continued to expand at an accelerated rate during the subsequent 72 hours and ultimately reached expansion levels comparable to those of continuously SIN-exposed cultures. These findings indicate that the effects of SIN stimulation are not solely dependent on continuous exposure to CCL21 and ICAM1 but rather reflect the induction of a durable biological program that remains active after withdrawal from the niche.

We further show that the persistence of this effect is not limited to proliferation. CAR T-cells transiently exposed to the SIN retained cytokine secretion and target cell killing capacity (Figure 4B) comparable to continuously SIN-treated cells.

Furthermore, morphological features associated with SIN exposure, included highly elevated cell size (determined by morphometric analysis of light microscopy images as well as by flow cytometry), elevated granularity, and polarized cellular morphology. Interestingly these morphological manifestations of the SIN-stimulated cells remained evident after transfer of the cells for further 3 days to non-coated plates (Figure 2). Together, these observations are consistent with the concept of sustained SIN-mediated cellular priming, which may be particularly relevant for clinical implementation because it raises the possibility that exposure of the CAR T-cells to the SIN for a limited time (10 days in the experiments reported here) might be sufficient to confer long lasting therapeutic benefits. Although not directly addressed in the present study, it is conceivable that such SIN-induced priming may persist after re-infusion of the cells into patients and contribute to an enhanced and long-term *in vivo* expansion, while retaining their antitumor activity. Future studies will be required to determine the duration and clinical relevance of this primed state following the adoptive transfer.

An additional consequence of the accelerated expansion of CAR T-cells is the potential for shortening of the manufacturing time. Clinically relevant CAR T-cell doses could potentially be achieved several days earlier than attainable with the current conventional culture methods, thereby reducing the duration of the *ex vivo* expansion. This may be particularly advantageous because prolonged culture is often associated with progressive differentiation, metabolic stress, replicative senescence, and decline or loss of cytotoxic efficacy. Consequently, SIN stimulation may enable the infusion of a “biologically younger” CAR T-cell product. Younger cellular products have repeatedly been associated with superior proliferative capacity, persistence, and higher therapeutic efficacy following adoptive transfer ^35,36^. Although these parameters were not directly evaluated in the present study, the accelerated manufacturing kinetics observed here suggest an additional potential translational benefit of SIN incorporation into CAR T-cell production workflows.

Beyond its effect on total cell yield, one of the most important findings of this study is the selective enrichment of the CAR-transduced sub-population of T-cells. As shown here, SIN stimulation had a remarkable and progressive effect on the prominence of CAR-transduced cells throughout the *ex vivo* expansion process, manifested by a persistent increase the frequency of CAR-positive cells from approximately 35–45% at early time points in culture to nearly 80% by day 13. Importantly, this increase was not merely a consequence of enhanced overall proliferation, but a clear preferential stimulation of the transduced cells by the SIN. Calculation of absolute cell numbers demonstrated a 14.6-fold increase in CAR T-cell expansion under continuous SIN exposure and a 19.4-fold increase following transient SIN exposure (Figure 3C, D). Interestingly, the frequency of CAR-positive cells remained essentially unchanged throughout culture under both standard non-coated conditions and full-scale GMP manufacturing without SIN exposure, highlighting the unique ability of the SIN to selectively enrich the transduced cell population.

The mechanism underlying the selective growth stimulation of the CAR-expressing cells is currently unclear. A possible explanation of this phenomenon, attributing this phenomenon to direct stimulatory effect of CCL21, via up-regulation of its receptor, CCR7, was ruled out since the levels of CCR7 on the CAR-expressing cells were low, compared to the CAR-negative cells. A more likely explanation is that CAR-expressing cells might exhibit a more sustained activation state that renders them more responsive to the combined chemokine (CCL21) and adhesion (ICAM1) signals. Consistent with this interpretation, SIN-treated cultures displayed increased expression of activation markers, including CD137 and CD69, together with progressive enrichment of the CAR-positive compartment, shown in Table 1.

Morphological and biophysical analyses further support the existence of a distinct SIN-induced activation program affecting CAR T-cells. SIN-CAR T-cells displayed enlarged cell size (e.g., larger projected area and a pronounced polarized morphology (low circularity) determined by light microscopy and higher Forward Scatter signal, and increased granularity, determined by Side Scatter values. These characteristics are consistent with cytoskeletal remodeling, metabolic activation, and active cellular proliferation ^24^. Similar morphological features were previously observed in SIN-treated TILs ^25^ and were associated with enhanced proliferative and cytotoxic potential.

From a translational perspective, the results presented here may be particularly valuable for the design of novel manufacturing platforms that utilize relatively low vector doses or exhibit variable transduction efficiencies, that could be compensated by the SIN’s capacity to increase the prominence of functionally active CAR T-cells by the preferential response of the CAR-transduced cells to the CCL21+ICAM1 SIN. Under such circumstances, increasing the absolute number and proportion of therapeutically active CAR T-cells without requiring additional viral vector input may improve manufacturing robustness, reduce production costs, and increase the probability of achieving clinically relevant cell yields within a shorter period of ex vivo expansion.

Finally, the enhanced nd sustained activation state of CAR T-cells, induced by the SIN was not accompanied by evidence of excessive terminal exhaustion. While expression of CD137 and CD69 was markedly increased, PD-1 expression remained relatively low and TIM-3 levels were not significantly altered by SIN exposure (Table 1). Moreover, SIN-expanded CAR T-cells retained robust antigen-specific IFN-γ secretion and efficient elimination of CD19-positive target cells. These findings suggest that SIN stimulation promotes productive activation without inducing overt functional impairment.

In conclusion, the novel findings reported here are consistent with a model in which a CCL21+ICAM1-based Synthetic Immune Niche reinforces specific signaling and adhesive cues, normally present within lymphoid tissues, thereby increasing the prominence of CAR T-cells with high therapeutic potency. We propose that incorporation of these lympho-mimetic signals into CAR T-cell production workflows may represent a simple and scalable strategy for improving both the yield and composition of CAR T-cell products.

## Acknowledgements

This study was supported by a grant (to BG) entitled: “Deciphering the principles of cell decision-making in multicellular systems: The Least micro-Environmental Uncertainty Principle (LEUP), awarded by the Volkswagen Foundation. The authors wish to thank Drs. Tamar Unger and Shira Albeck of the Dana and Yossie Hollander Center for Structural Proteomics (WIS) for the expert production of the SIN components used in this study. We also thank Yosef Addadi and Ofra Golani of the de Picciotto Cancer Cell Observatory (WIS) and the Flow Cytometry Unit (WIS) for their expert assistance.

Furthermore, the authors would like to thank the Samueli Foundation for their generous support of this study (to MJB).

