## Supplemental Method and Figures for "A Lymphomimetic Synthetic Immune Niche, Consisting of CCL21 and ICAM1, Accelerates the Expansion of Potent CAR T-cells"

**Supplementary Methods**

Production and purification of human CCl21 (hCCL21):

The gene encoding hCCL21 (24-134) followed by a TEV cleavage site, and a C-terminal 6 x His tag was cloned into pET21a. The resulting construct was transformed into E. coli Bl21 (DE3) competent cells. Bacterial pre-cultures were grown overnight at 37°C with shaking and used to inoculate 7.5L LB. Cultures were grown at 37°C with shaking to an optical density OD600= 0.6. Protein expression was induced by the addition of isopropyl-β-d-thiogalactopyranoside (IPTG; 200 μM), and the culture was kept at 15°C with shaking overnight. Cells were harvested by centrifugation, and the pellet was resuspended in 100 ml lysis buffer containing 50mM Tris pH 8, 300mM NaCl,10mM Imidazole,1mM DTT supplemented with 0.2mg/ml lysozyme, 20ug/ml DNase, 1mM MgCl2, and protease inhibitor cocktail (Calbiochem set 3). After lysis by a cooled protein disrupter (Constant Systems), the soluble fraction was obtained by centrifugation and purified by immobilized metal ion affinity chromatography (IMAC) using a HiTrap FF5ml cartridge (Cytiva) using an FPLC system (ÄKTA GE Healthcare Life Sciences) at 4oC. The column was washed with lysis buffer containing 50mM imidazole and hCCl21 was eluted in one step with elution buffer (50mM Tris pH 8, 300mM NaCl, 500mM imidazole). The fractions containing hCCl21 were analyzed using SDS-PAGE, pooled and dialyzed into PBS overnight. hCCl21 was applied to an Xbridge C4 (Waters) reverse phase column operated by an HPLC system (Waters) pre-equilibrated with binding buffer containing 5% acetonitrille, 0.1%TFA in DDW. The protein was eluted with a linear gradient to 95% acetonitrille, 0.1%TFA in DDW. Pure hCCl21 samples from consecutive runs were pooled and lyophilize to dryness. The dry protein was resuspended in DDW and divided into small aliquots before flash freezing in liquid nitrogen.

Expression and purification of human ICAM1 (hICAM1):

hICAM1 (aa residues 28-480) followed by an Fc domain, a TEV recognition site, and a C-terminal 6 x His tag was cloned into pHLSec vector for secretion from Expi293 human cells. Medium (300ml) containing the secreted protein was collected 72 hr. post transfection for purification. After carefully removing the cells, the medium was applied to a Ni column (HisTrap_FF_5ml) equilibrated with PBS and the bound hICAM was washed with PBS containing 50mM imidazole and eluted with PBS containing 500mM imidazole. Fractions containing hICAM1 were pooled, concentrated and injected to a Superdex_200_16/60_PrepGrad size exclusion column equilibrated with PBS. The identity of the protein was confirmed by SDS PAGE developed with Coomassie blue. Aliquots were flash frozen with liquid nitrogen and kept at -80◦C.

### Supplementary Figures

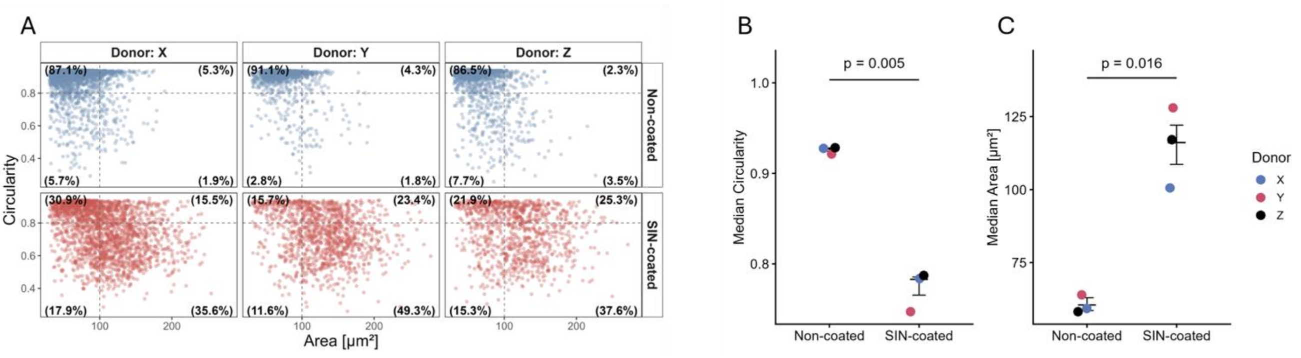


**Figure S1: CCL21+ICAM1 treatment increases the prominence of large and polarized (low circularity) CAR T cells.** CAR T cells were cultured on SIN-coated or non-coated plates for 13 days and imaged on day 9. **(A)** Quantification of projected cell area and circularity of SIN-treated and untreated CAR T cells. A projected area threshold of 100 µm² was used to distinguish small from large cells, and a circularity threshold of 0.8 was used to distinguish circular from polarized cells. **(B-C)** Median CAR T-cell circularity **(B)** and projected area **(C)** are shown for each donor and condition. P-values were calculated using paired t-test.

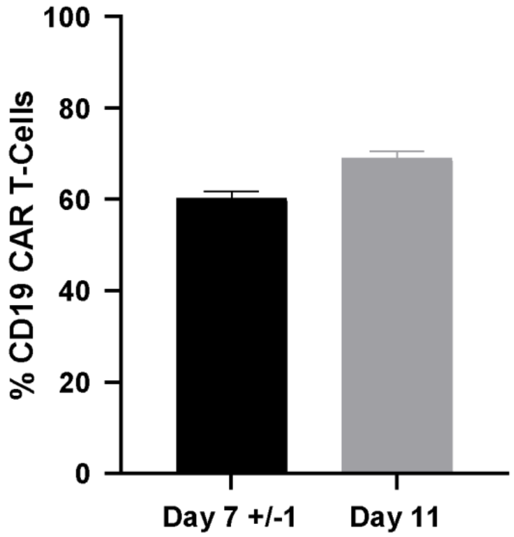


**Figure S2. CAR T-cell transduction efficiency during GMP manufacturing**. The percentage of CAR-positive T cells was assessed on day 7 ± 1 and day 11 of culture. Data represent mean ± SD (n=4).
